# Estrogen regulates early embryonic development of the olfactory sensory system via estrogen-responsive glia

**DOI:** 10.1101/2021.06.03.446864

**Authors:** Aya Takesono, Paula Schirrmacher, Aaron Scott, Jon M. Green, Okhyun Lee, Matthew J. Winter, Tetsuhiro Kudoh, Charles R. Tyler

**Affiliations:** Biosciences, College of Life and Environmental Sciences, University of Exeter, Exeter, Devon, EX4 4QD, United Kingdom; Department of Biological and Marine Sciences, Faculty of Science and Engineering, University of Hull, Hull, HU6 7RX, United Kingdom; School of Physiology, Pharmacology & Neuroscience, University of Bristol, Bristol, BS8 1TD, United Kingdom

**Keywords:** Endocrine disrupting chemicals, Transgenic zebrafish embryo, embryogenesis, Olfactory glomerular development, Radial glia progenitor cells, Olfactory-mediated behaviour

## Abstract

Estrogen is well-known to regulate development of sexual dimorphisms of the brain, however its role in the brain during early embryonic development prior to sex-differentiation is unclear.

Using estrogen biosensor zebrafish models, we found that estrogen activity in the embryonic brain occurs specifically in a type of glia located within the OB, which we name estrogen-responsive olfactory bulb/EROB cells. With estrogen activity, EROB cells extend their ramified projections that overlay the OB outermost layer and tightly interact with olfactory sensory neurons (OSNs) at the olfactory glomeruli. Pharmacologically inhibiting estrogen activity and/or EROB cell ablation impedes olfactory glomerular development, including OSN pathfinding, topological organisation of olfactory glomeruli and inhibitory neurogenesis in the OB. Furthermore, activation of this estrogen/EROB-dependent mechanism decreases the intrinsic neuronal activity primarily in the OB, and this alteration of estrogen signalling disrupts olfaction-mediated behaviour. We propose that estrogen acts on glia to regulate development of functional OB circuits, thereby modulating the local intrinsic excitability in the OB and olfaction-mediated behaviour. Our data also suggest a possibility that the estrogen/EROB cascade may be an important site of action for environmental estrogens causative of neurodevelopmental impairments in animals and humans.

## Introduction

Estrogen is a signalling molecule that plays critical roles in the development of reproductive and somatic organs, including the brain. In vertebrates, estrogen is supplied to embryos maternally through the placenta, or in oviparous species through yolk provisioned into eggs. Estrogen can also be produced locally within the embryonic brain through the conversion of testosterone to estrogen by the enzyme, aromatase (Bondesson et al., 2015; Menuet et al., 2005). In rodents, it is thought that the embryonic brain has greater estrogen activity compared with the brain in later life stages, as evidenced by the fact that the highest levels of estradiol (the most potent endogenous estrogen), aromatase activity, and expression of estrogen receptors (two isoforms of nuclear receptors, ERα and ERβ) occur in the brain at this time (George and Ojeda, 1982; Konkle and McCarthy, 2011; McCarthy, 2008). The importance of estrogen signalling pathways in the developing brain has been further supported by neurodevelopmental phenotypes in estrogen receptor (ER), ERα and ERβ, knockout (KO) mice. Studies on ERαKO mice have shown that ERα contributes to the establishment of the hypothalamic–pituitary–gonadal axis and is required for sexually dimorphic circuitry and sex-specific behaviour (Ogawa et al., 1998a; Ogawa et al., 1998b; Simerly et al., 1997; Stephens et al., 2016); In contrast, ERβ^-/-^ mice show marked neuronal loss in somatosensory cortex and defects in the development of calretinin^+^ GABAergic interneurons in specific regions of the brain (i.e. cortex, hypothalamus and amygdala) in the prenatal period (Fan et al., 2006; Wang et al., 2001; Wang et al., 2003), which may link with altered spatial learning and behaviours seen in ERβ^-/-^ adult mice (Liu et al., 2008; Rissman et al., 2002). ERβ is also critically involved in neural specification of pluripotent stem cells in vitro (Varshney et al., 2017). ER-mediated transcriptional activation occurs mainly in the neural ectodermal tissues from E12 in the estrogen responsive element (ERE)-Luciferase mouse model (Della Torre et al., 2018) further evidencing the importance of estrogen signalling in brain development.

However, there is little understanding on how estrogen exerts its effects in the embryonic brain before the period of sex differentiation, what the physiological effects of estrogen are, or what the developmental consequences are for alterations to estrogen signalling in the brain during embryogenesis. This understanding has a further level of importance given that early-life exposure to environmental contaminants – so-called endocrine disrupting chemicals (EDCs) - that mimic estrogens is thought to cause a wide range of adverse impacts on brain development and function (Derouiche et al., 2015; Kinch et al., 2015; Lichtensteiger et al., 2015; Porseryd et al., 2017) which in turn may lead to defects in cognition, learning, emotional control and behaviour in later life (Braun et al., 2009; Derouiche et al., 2015; Gioiosa et al., 2013; Kawai et al., 2003; Kinch et al., 2015). In fact, emerging evidence suggests that EDC pollutants in the environment have impacted natural habitats and affected social and reproductive behaviours in wildlife populations (Godfray et al., 2019; Nash et al., 2004; Soffker and Tyler, 2012). EDC-induced impairment of estrogen functions during brain development have also been implicated in the pathogenesis of some sex-biased neurodevelopmental diseases in humans, including attention-deficit hyperactivity disorder (ADHD), autism spectrum disorders and schizophrenia (Crider and Pillai, 2017; Mustieles et al., 2015). Thus, a suitable model to study estrogen’s contributions to embryonic brain development in real-time has been highly sought after.

In this work, combining the use of estrogen biosensor (Green et al., 2016; Lee et al., 2012a), calcium sensor (Winter et al.; Winter et al., 2017) and chemical/genetic cell ablation zebrafish models, we reveal a new physiological function of estrogen in the early developing brain which commences shortly after the initiation of neurogenesis. Our findings demonstrate for the first time that estrogen exerts a highly specific effect on the olfactory bulb (OB) of the embryonic brain and critically contributes to the development and function of olfactory sensory system through newly identified target glia. Given that this occurs shortly after the regional specification of the embryonic brain, at a much earlier stage than the initiation of sex differentiation in zebrafish (i.e. 20-25 dpf) (Lau et al., 2016; Uchida et al., 2002), this estrogen-mediated cascade is likely a fundamental mechanism required for development of olfactory sensory system, regardless of sex.

## Results

### The estrogen/ER-mediated transcriptional activation occurs in the olfactory bulb in the zebrafish embryonic brain

To understand the role of estrogen in embryonic brain development, we used an estrogen biosensor zebrafish model (ERE:GFP) which allows the identification of *in vivo* cell responses through estrogen/ER-mediated induction of GFP (Green et al., 2016; Lee et al., 2012a; Lee et al., 2012b). When ERE:GFP embryos were exposed to ethinylestradiol (EE2, 100 ng/L), a synthetic derivative of the female sex hormone 17β- estradiol, we observed that estrogen response (indicated by GFP expression) occurred primarily in the olfactory bulb (OB) located in the anterior-most region of the forebrain (Fig. 1Aii, 48 hours post fertilisation (hpf)), where olfactory signals are first processed in the brain (Miyasaka et al., 2013). These GFP-positive cells were located around the midline ventricular region of the anterior forebrain. By 96 hpf the number of these estrogen responsive cells in the OB (which we name EROB cells) was much increased (Fig. 1B and 1Ci) with their projections extending laterally towards the pia of the OB (Fig. 1Ci-ii). This unique morphology and location of EROB cells were further confirmed by injecting UAS-DsRed DNA into ERE:GFP embryos, which renders a mosaic expression of DsRed in a subset of the EROB cells (Fig. 1Di-ii). We next investigated the ontogeny of the EROB cells during embryonic and larval brain development. We detected the EROB cells with the cellular projections starting to develop after 32 hpf (Fig. 1E) and with complex projection networks formed by 48hpf (Fig. 1F). The number of EROB cells greatly increased between 48hpf and 72hpf (Fig. 1G), reaching a maximum number at around 4-5 dpf (Fig. 1J) and then gradually decreased thereafter (Fig. 1H, 1I and 1J). In later life stage (15 and 21dpf), the size (length) and number of EROB cell projections were seen to be markedly reduced and the somata now dispersed from their original position at mediodorsal OB (Fig. 1H and 1I). The localisation of the EROB cells and their ontogenic profiles suggest they have a role in olfactory development in early embryo-larval brain development.

**Figure 1.**
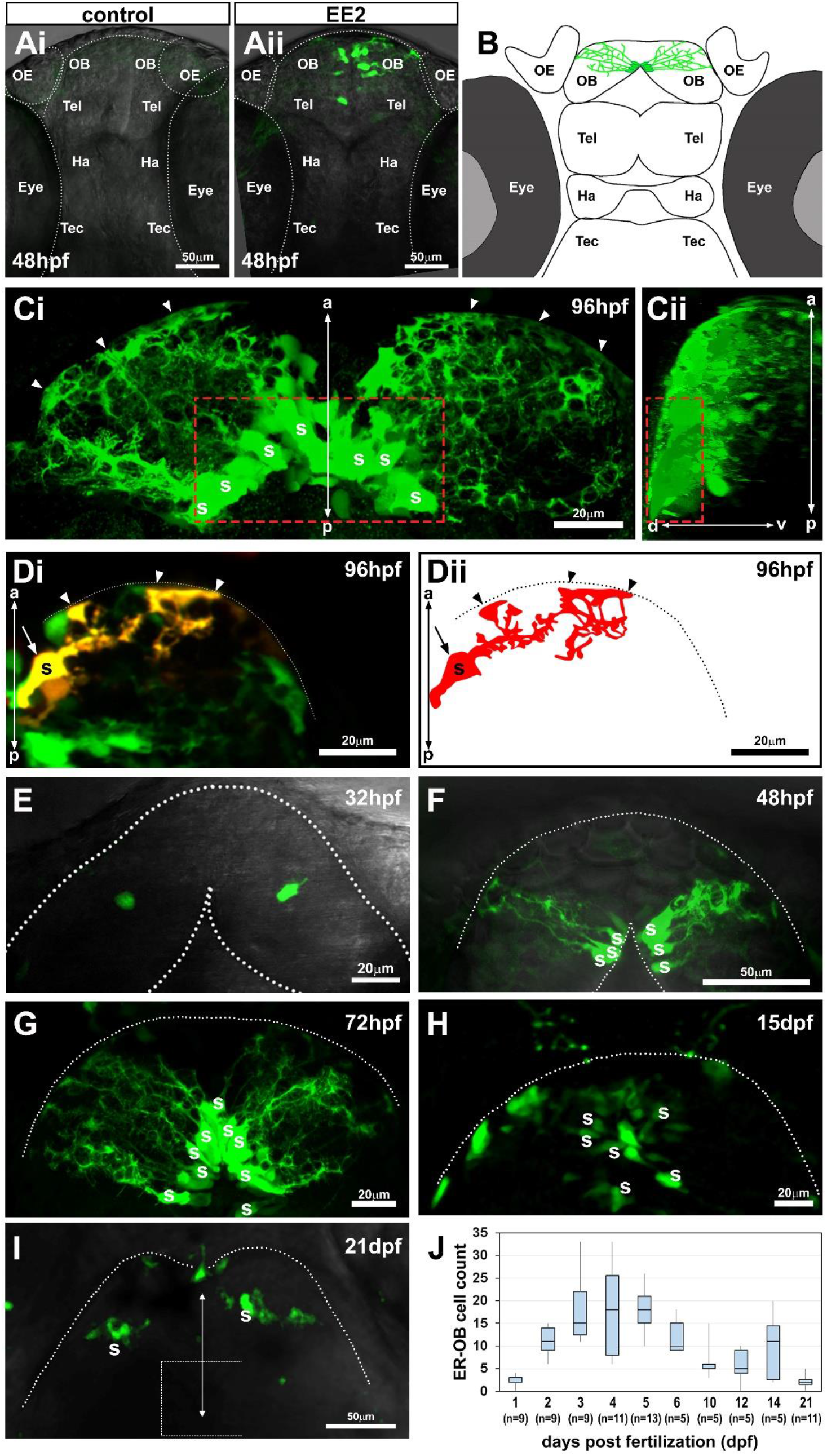
The earliest estrogen/ER-mediated transcriptional activation occurs in a small number of cells in the olfactory bulb in the zebrafish embryonic brain. (Ai-ii) Confocal z-projection images of control (Ai) or 17-α ethinylestradiol (EE2, 100 ng/L, Aii) - exposed zebrafish ERE:GFP embryos at 48 hpf. OE, olfactory epithelia; OB, olfactory bulb; Tel, telencephalon; Ha, habenula; Tec, tectum; Eye. (B) Illustration of the Estrogen Responsive Olfactory Bulb (EROB) cellular domain in a 4 dpf zebrafish embryo. (Ci-ii) Confocal z-projection images (Ci, dorsal view; Cii, a 90° rotated view) of EROB cells in EE2-exposed ERE:GFP embryos at 96 hpf. Red dotted rectangle indicates the mediodorsal OB. Some of the somata are marked as “s”. Arrow heads, indicate the distal ends of EROB cells; anterior-posterior (a-p) axis; dorso-ventral (d-v) axis. (Di-ii) Di shows the morphology of a single EROB cell from a confocal z-projection image and Dii illustrates a trace of the cell morphology. The midline, left edge; Dotted lines, the OB pia; Arrow heads, the termini of EROB cells at the OB pia. The somata (marked “s”) of the EROB cell is indicated with the arrow. (E-I) Confocal z-projection images of EROB cells (green) showing their ontological development in the brain of embryo - larval zebrafish (dorsal view). Dotted lines, the OB pia; “s” indicates the EROB cell somata; a white dotted square in (I), the mediodorsal OB. (J) Ontogenic profiles of EROB cell numbers.

### EROB cells are aromatase B positive glia

The morphology of the EROB cells and their location in the OB suggest they may be glial cells. In fact, we found that the projections from these EROB cells labelled strongly with anti-Glial Fibrillary Acidic Protein (GFAP) antibody, a marker for astrocytic glial cells (Fig. 2Ai-ii). The EROB cells also expressed a brain specific estrogen synthesising enzyme, aromatase B gene (*cyp19a1b*) as shown by their overlap with *cyp19a1b:GFP* (Brion et al., 2012) in double transgenic embryo carrying both *ERE:mCherry* and *cyp19a1b:GFP* reporter genes (Fig. 2Bi-ii). Co-localisation of EROB cell and GFAP in the same projection (Fig. 2Aiii) and co-localisation of EROB cells and aromatase B expression (Fig. 2Biii) were confirmed by the application of method based on Pearson’s correlation coefficient in two dimensions (Steinfeld et al., 2015). We also found that EROB cells co-expressed Sox2 (a maker of neuronal progenitor cells) (Fig. S1Ai-iv, Fig. S1Bi-ii) but they were segregated from the brain region where elavl3/HuC-positive immature post-mitotic neurons were localised (Fig. S1Ci-iv, Fig. S1Di-ii). These data demonstrate that EROB cells are not neurons but Sox2^+^/GFAP^+^/aromatase B^+^ expressing glia.

**Figure 2.**
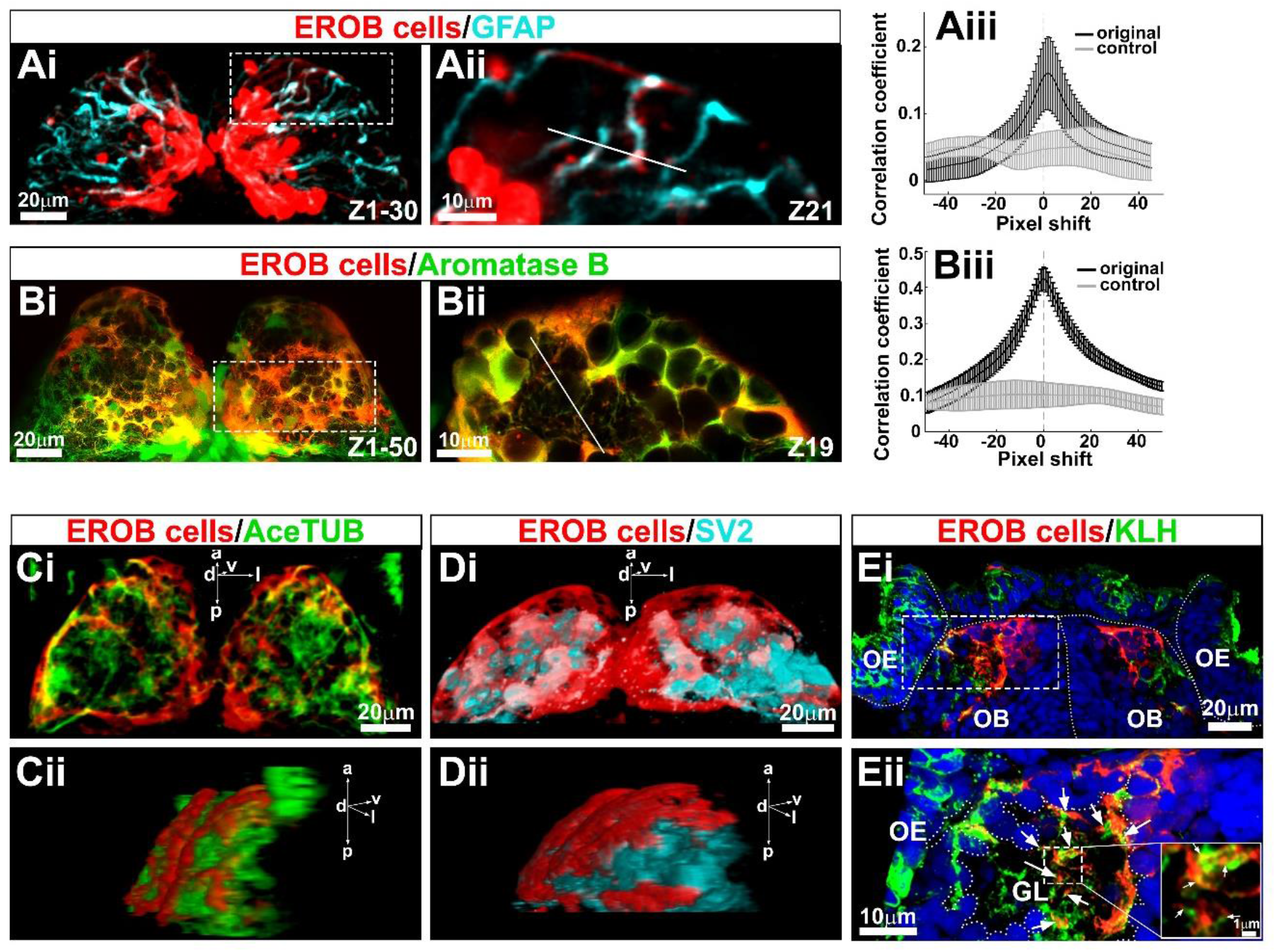
Estrogen Responsive Olfactory Bulb (EROB) cells are GFAP and aromatase B expressing glia that interact with olfactory sensory neurons at olfactory glomeruli. (Ai) Confocal z projection images of EROB cells (mCherry, red) and GFAP (cyan) in an EE2-exposed 4 dpf ERE:mCherry embryo. (Aii) An optical section image (1.5 μm step size) magnified from the area marked with white dotted rectangle in Ai. (Aiii) Pearson’s correlation coefficient in two dimensions for EROB cells vs. GFAP which was plotted against image displacement in the x direction (black). Control corresponds to image sets with one channel rotated by 90° (light grey). The peak correlation coefficient (PCC) was found at a 1.5 pixel shift (0.22μm/pixel size), suggesting a close localisation of EROB cell and GFAP in the same projection. (Mean values ± SEM shown). (Bi) Confocal z projection images of 4 dpf EE2-exposed double transgenic - Tg(*ERE:mCherry*) x Tg(*cyp19a1b:GFP*) - embryo. (Bii) One optical section image (0.7 μm step size) magnified from the area marked with white dotted rectangle in Bi. (Biii) Pearson’s correlation coefficient in two dimension for EROB cells vs. aromatase B. The PCC was at 0 pixel shift, indicating co-localisation of EROB cell and aromatase B. p* < 0.05, t-test for significant difference from the rotated control image sets (grey). Mean ± SEM shown (n=3). (Ci-ii) A 3D image of EROB cells (red) and acetylated tubulin positive axonal projections (green) in the OB (Ci, dorsal view; Cii, a 50° rotated view of Ci). d, dorsal; v, ventral; l, lateral; a, anterior; p, posterior. (Di-ii) A 3D image of EROB cells (red) and olfactory glomeruli stained with SV2 antibody (cyan) (Di, dorsal view; Dii, a 45° rotated view of Di). (Ei-ii) Frontal cryosection image of EROB cells (red), OSNs (KLH, green) and nuclei (Hoechst, blue). White dotted lines indicate the outline of the olfactory epithelia (OE), olfactory bulb (OB) and midline. (Eii) A magnified image of a glomerulus (GL) from the area indicated with a white dotted rectangle in Ei. Arrows indicate EROB cell - KLH^+^ OSN interactions. Outline of the GL is indicated with white dotted freeform line. Magnified image of a central area of the GL is inserted on the right, showing that EROB cells (red) tightly intermingle with OSNs (green) within the GL (indicated with white arrows).

### EROB cells interact with olfactory sensory neurons at olfactory glomeruli via their projections

We next examined for connectivity between EROB cells and neurons in the OB by co-staining EROB cells (mCherry) with antibodies against anti-acetylated tubulin, a marker for axonal projections, and against anti-synaptic vesicle glycoprotein 2 (SV2), a marker for synaptic neuropils, including olfactory glomeruli. We confirmed that the projections of the EROB cells spanned the entire surface of the OB (Fig. 2Ci-ii, Suppl. Movie 1) and overlaid olfactory glomeruli (Fig.2Di-ii, Suppl. Movie 2). The distal ends of EROB cell projections were associated closely with the acetylated tubulin^+^ axons at olfactory glomeruli (Fig. S2A, Bi-iv) and surrounded each SV2^+^ glomerulus (Fig. S2C, Di-iv). The interaction between EROB cells and olfactory sensory neurons (OSNs) were furthermore examined by co-labelling EROB cells (mCherry) with an antibody against anti-keyhole limpet haemocyanin (KLH), a specific maker for OSNs, in frontal cryosections of the OB. This showed that the EROB cells projected to the glomerulus at the point where they connected with the axonal termini of KLH^+^ OSNs (Fig. 2Ei-ii). These projections from EROB cells intricately intertwined with KLH^+^ OSNs in the olfactory glomeruli (Fig. 2Eii, inserted high magnification image). These data suggest that EROB cells may act as cellular guides for directing the axonal growth of OSNs to specific glomeruli in the OB and for winding the termini of OSNs into a spherical shape within a glomerulus.

### Estrogen regulates EROB cell projections

We then questioned whether ER-mediated activation of estrogen signalling is required for the establishment of EROB cell projections, which appear to be an integral structural component of the OB. To address this, ERE:GFP embryos were treated with EE2 from 1 hpf to 72 hpf to induce GFP in EROB cells and then underwent either a washout period between 72hpf and 120hpf (referred as “control”) or subsequent treatment with an ER antagonist (ICI182,780, ICI) to inhibit estrogen activity between 72hpf to 120hpf (“ICI group”). The control group showed intense EROB cell projections overlaying the majority of OB surface (Fig. 3Ai, Fig. 3Aii). In contrast, the ICI treated group showed substantial alteration to the EROB cell morphology (Fig. 3Bi, Fig. 3Bii). To quantify this effect of the ICI, mean GFP intensity within the EROB cell margin (marked with a green square in Fig. 3Ai or 3Bi) was plotted along the width of the OB margin. The ICI treated group showed a consistent reduction in GFP signals near the OB pia compared with the control group, representing distorted EROB projections in ICI groups (Fig.3C; Fig. 3E and 3F, p** < 0.01, Mann Whitney U test). This ICI effect was unlikely to be due to an effect on EROB cell number as the EROB cell count did not differ between the control vs. treatment group (Fig. 3G, p = 0.19, t-test). These data indicate that estrogen activity is essential for the establishment of EROB cell projection networks during OB development.

**Figure 3.**
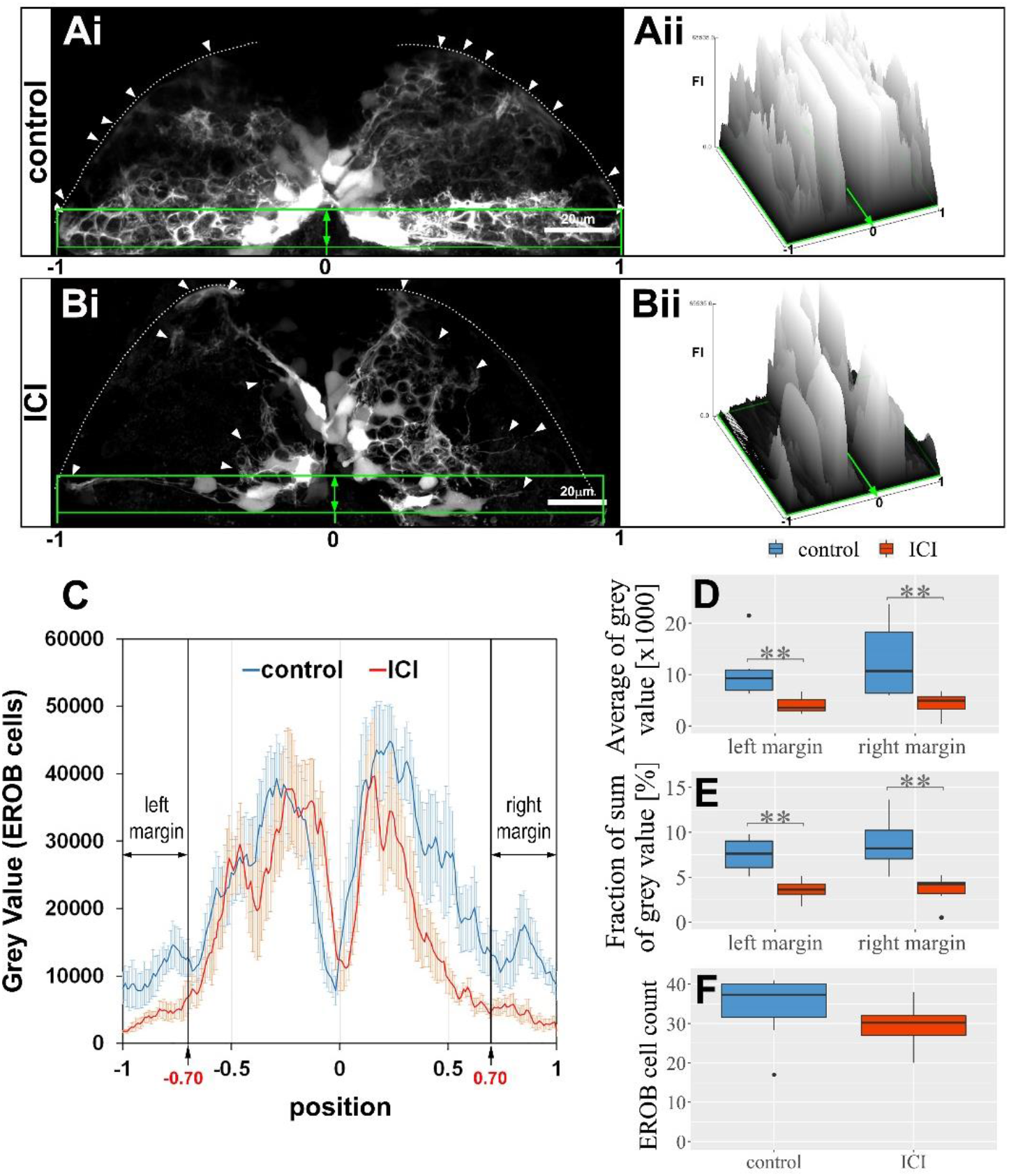
Estrogen regulates EROB cell projections. (Ai and Bi) Confocal z projection image of EROB cells in EE2-exposed control (Ai) or subsequently an estrogen receptor antagonist, ICI 182,780 (ICI, 10 μM)-treated (Bi) 5dpf ERE:GFP embryo (see details in “Methods”). Arrow heads, the distal end of the EROB cells; white dotted line, the pia of the OB; green rectangles, EROB cell margins. (Aii and Bii) Surface plots of FI of GFP signals (EROB cells) in the EROB cell margins in control (Ai) and ICI-treated (Bi) embryos. The surface plots are shown along the width of the EROB cell margin, adjusted by the position of the midline as “0” and the positions of both ends of the margin as “-1 (left)” or “1 (right)”. (C) The intensity plots of GFP signals (EROB cells) in the EROB cell margin in control (blue) and ICI (red) embryos. X-axis is adjusted as described above. Mean ± SEM shown, N=6 each. Black both-end arrows indicate both edges of the EROB cell margin, at the positions of [−1:−0.7](“left margin”) and [0.7:1](“right margin”). (D and E) The average of grey value and sum of grey value (% of total grey value in each embryo) of left and right margin (N = 6 each, p** < 0.01, Mann Whitney U test). (F) The EROB cell number (N = 8).

### Estrogen/EROB cells are essential for olfactory glomerular development

To assess the inter-relationship between estrogen and the EROB cells in the development of the OB we blocked the estrogen signalling through the application of ICI, or ablated the EROB cells using nitroreductase-mediated chemical/genetic cell ablation (Curado et al., 2008) (details in “Methods”; Fig. S3A and 3B). In this work we measured the locations and volumes of five different olfactory glomeruli - medioanterior glomerulus (maG), dorsal glomerulus (dG), dorsolateral glomerulus (dlG) and mediodorsal glomerulus 3 (mdG3) and a group of mediodorsal glomeruli (mdG1-6). These were selected because they are located in close proximity to the dorsal surface of the OB (Braubach et al., 2013) and thus likely to be interacting directly with EROB cells. In addition, we examined the axonal extension patterns of OSNs projecting towards olfactory glomeruli. EROB cell ablation (EE2 + metronidazole (MTZ)) and pharmacological inhibition of estrogen signalling (via ICI) altered the stereotypical positions and volumes of some of OB glomeruli: with EE2+MTZ or ICI, mdG1-6 were often ectopically positioned, for instance, being closer to the midline as compared with the control condition (Fig. 4A, SV2 images and illustration of OB glomeruli map). In addition, mdG3 was much smaller in size or was absent in EE2+MTZ and ICI exposed embryos (Fig. 4A). Of the five examined glomeruli, maG and mdG3 were the most severely affected by the EE2+MTZ or ICI treatments, leading to reductions in their volumes (Fig. 4B, maG and mdG3, p* < 0.05, ANOVA with Tukey’s test). However, ICI causes broader effects compared with EE2+MTZ (Fig. 4B, dG and dlG, p* < 0.05, ANOVA with Tukey’s test). We also observed atypical axonal projections of OSNs in EE2+MTZ and ICI treated embryos with their projections now extended towards the midline of the OB (Fig. 4A, yellow arrows in KLH images). These phenotypes were not seen in control samples or embryos treated with EE2. These data support that estrogen and EROB cells are critically involved in olfactory glomerular development.

**Figure 4.**
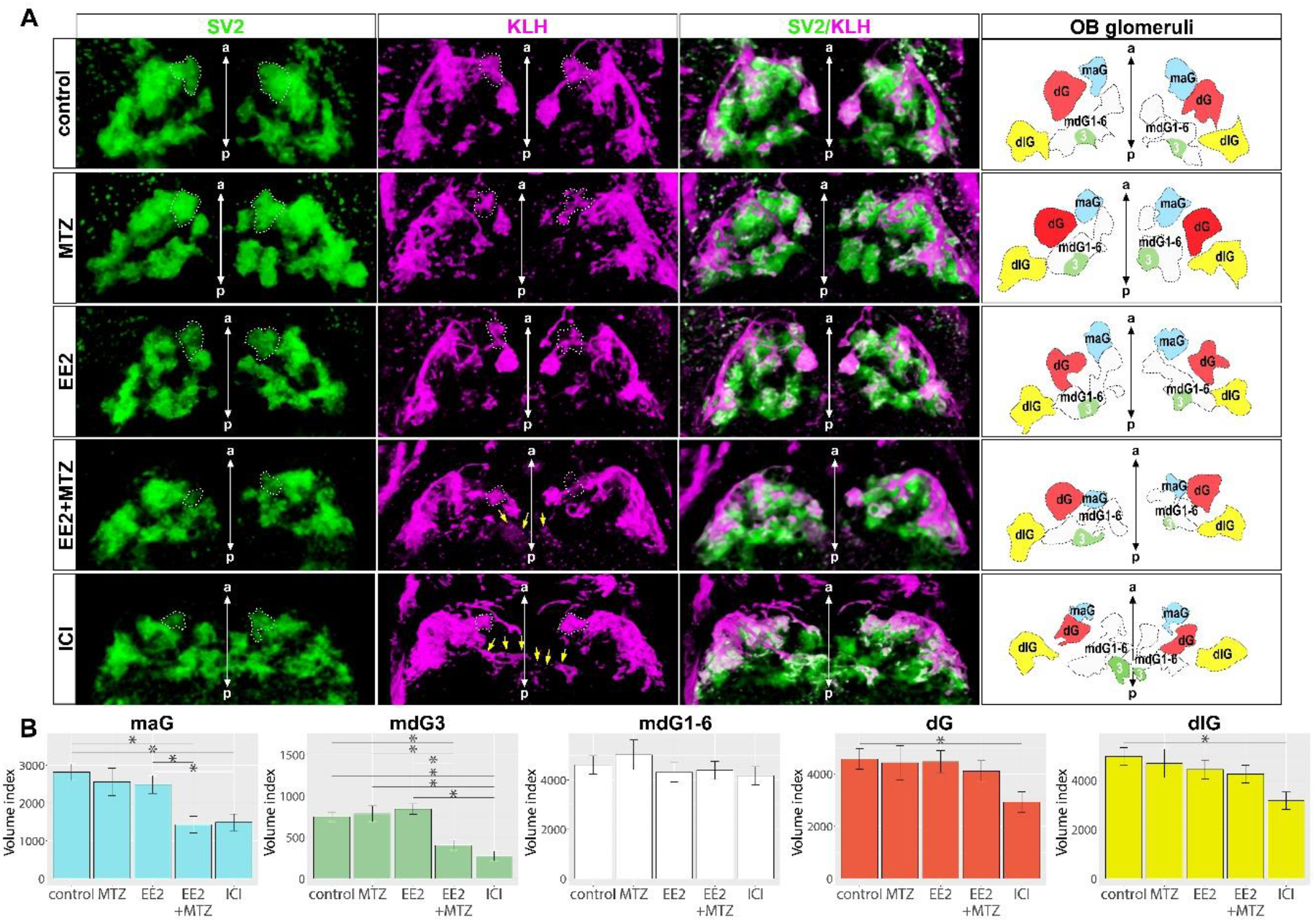
EROB cell ablation impairs the development of olfactory glomeruli. (A) Representative 3D images of olfactory glomeruli (SV2, green), OSNs (KLH, magenta) and merged image of SV2/KLM in control, MTZ, EE2, EE2+MTZ or ICI treated 4 dpf ERE:mCherry embryos. Right-end panels are the illustrations of OB glomeruli map generated based on the outlines and positions of 5 selected olfactory glomeruli in the representative 3D images: maG, blue; dG, red; dlG, yellow; mdG1-6, white and mdG3, green. The midlines are indicated with a-p axis in the images. White dotted lines in SV2 and KLH images, maG; green coloured areas in OB glomeruli map, mdG3; yellow arrows in KLH images, altered axonal extensions of OSNs. (B) Relative volumes of the 5 different selected glomeruli. N=8 for control, EE2, EE2+MTZ and ICI, N=3 for MTZ alone, Mean ± SEM shown. p* < 0.05, ANOVA with Tukey post-hoc test.

### Estrogen promotes inhibitory synapse formation in the OB

Our findings show that estrogen signalling and its target EROB cells play a key instructive role in olfactory glomerular development. It is known that inhibitory regulatory circuits at olfactory glomeruli, which comprise of various types of inhibitory interneurons, are consecutively established during the course of olfactory glomerular development (Batista-Brito et al., 2008; Nagayama et al., 2014). We therefore examined if estrogen signalling and EROB cells could also regulate the local synapse formation in the OB. To address this, firstly we co-stained EROB cells (GFP) with gephyrin, a postsynaptic scaffolding molecule for inhibitory synapses (Tyagarajan and Fritschy, 2014) to analyse the spatial relationship between EROB cells and gephyrin+ inhibitory synapses in the OB. As shown in Fig. 5Ai-ii, EROB cell projections were located closely with gephyrin expressing puncta at olfactory glomeruli. We found that the EE2 treatment increased the number of gephyrin+ puncta in the OB (Fig.5Ci-ii) whereas inhibiting estrogen signalling by ICI markedly reduced the number of the puncta (Fig.5Di-ii) as compared with those in control (Fig.5Bi-ii). Notably EE2 increased both the number of gephyrin+ puncta and also the puncta size, suggesting EE2 may promote clustering of gephyrin proteins. This effect of EE2 was greater in the larger sized puncta (Fig. 5E, 1-10μm^2^; control vs EE2, p* < 0.05 ANOVA with Tukey’s test; 0.1-0.4μm^2^; control vs EE2, p = 0.18 ANOVA with Tukey’s test). Conversely, when estrogen activity was inhibited by ICI, the number of gephyrin puncta was greatly reduced (Fig. 5E), indicating that estrogen signalling is essential for establishing inhibitory synapses in the OB. These data suggest that estrogen regulates inhibitory synaptogenesis.

**Figure 5.**
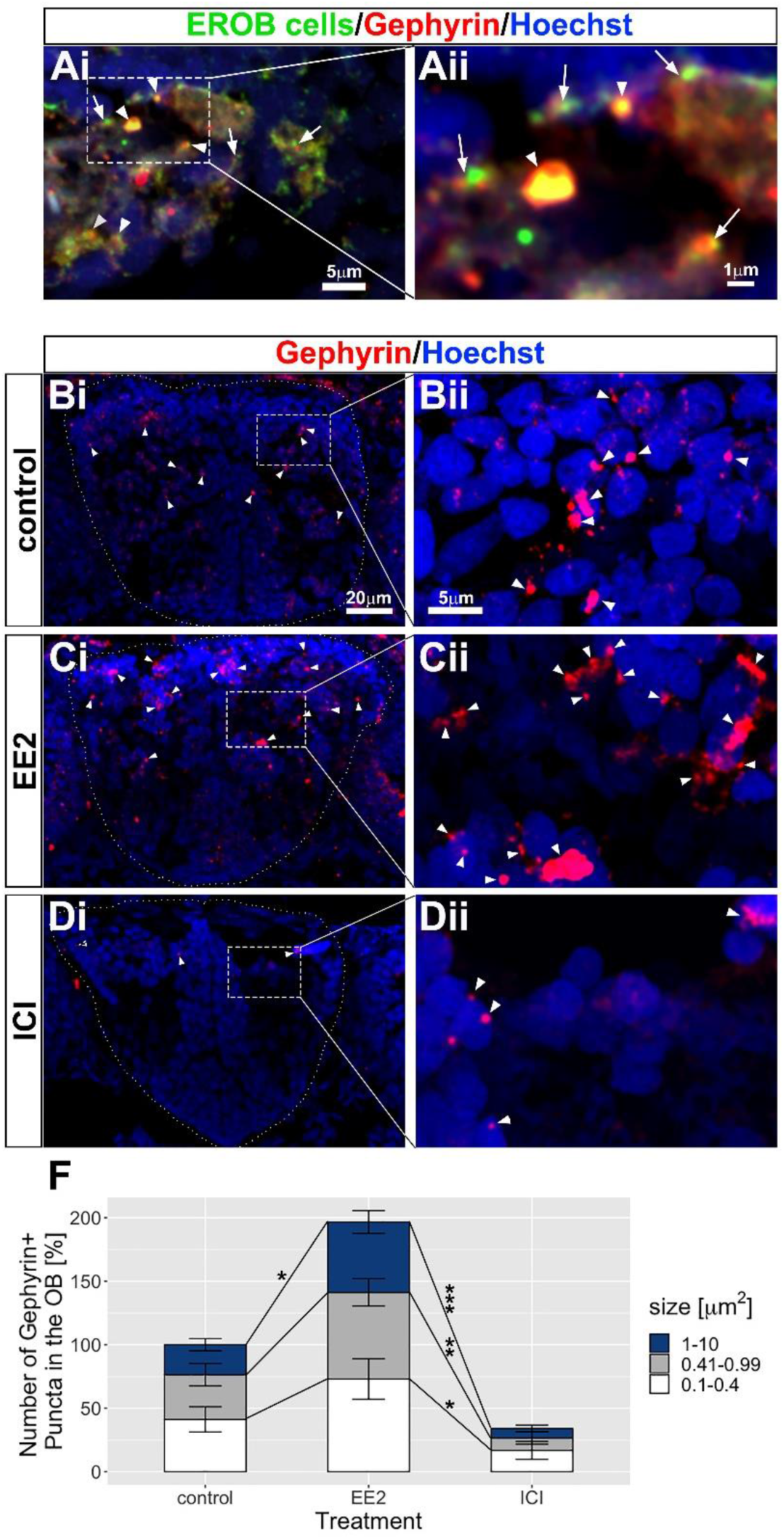
Estrogen promotes inhibitory synaptogenesis in the OB. (Ai-ii) A transverse cryosection image of EROB cells (green), gephyrin puncta (red) and nuclei (blue). The distal-ends of EROB cell projections co-localise with (arrow heads) or localise in close proximity to (arrows) gephyrin+ inhibitory synapses at an olfactory glomerulus. (Aii) A magnified image of a sub-region of olfactory glomerulus in Ai. (Bi-ii, Ci-ii and Di-ii) Estrogen promotes inhibitory synaptogenesis in the OB while ICI inhibits the formation of inhibitory synapses. (Bi-ii), control; (Ci-ii), EE2-treated; (Di-ii), ICI-treated 4 dpf ERE:GFP embryos. Gephyrin puncta (red) in the OB are shown with arrow heads. (Bii, Cii and Dii) Magnified images of a sub-region of the OB indicated in Bi, Ci and Dii. (F) % of increase in the total numbers of gephyrin^+^ puncta in the OB. The number of gephyrin^+^ puncta was quantified for three different sizes, 0.1-0.4 μm^2^(white), 0.41-0.99 μm^2^(grey) and 1-10 μm^2^(blue). Mean ± SEM shown. N=4. p* < 0.05; p** < 0.01; p*** < 0.001. ANOVA with Tukey post-hoc test.

### Estrogen signalling and EROB cells specifically modulate the intrinsic/spontaneous excitability in the OB in the embryonic brain

Given that estrogen/EROB cell cascade critically and selectively regulates olfactory glomerular development, we examined if alteration of the estrogen signalling affects local neuronal activity in the OB in the developing embryos. To do that, we employed a calcium sensor *elavl3/HuC:GCaMP6s* transgenic zebrafish model in combination with light sheet microscopy (LSM) and a GCaMP6s image processing pipeline established previously in our laboratory (details in “Methods”) (Winter et al., 2017). Using this system, a full brain volume of GCaMP6s images can be captured that allows us to extract region of interest (ROI)-specific neuronal activity data. We found that developmental exposure to EE2 (from 1 hpf to 4dpf) most significantly reduced the intrinsic neuronal activity in the olfactory bulb (OB) and OB glomerular layer (OBGL) out of 41 different ROIs compared with those in control condition (Fig. 6A; and Fig. 6B, marked with black rectangle). Conversely, developmental exposure to ICI specifically increased GCaMP signals in the OB and the OBGL. (Fig. 6A, ICI, highlighted in red; Fig. 6B). Dose-response experiments confirmed that the effects of EE2 and ICI on the intrinsic excitability were inverse, dose-dependent and specifically observed in the OB and OBGL but not in other brain regions, such as cerebellum (Fig. 6C). To clarify whether this OB specific change in excitability was due to the acute response to the estrogen signalling pathway (Kramar et al., 2009; Mukherjee et al., 2017; Zhang et al., 2010) or in fact a long-term response, we examined the effect of transient exposure to EE2 or ICI on the intrinsic neuronal activity in the same assay system. Unlike the developmental exposure to EE2, transient EE2 exposure (20 minutes before recording) did not cause changes in excitability in the OB (Fig. S4). Furthermore, the impacts of transient exposures of ICI were not specific to the OB and were more widespread (albeit more mild responses, Fig. S4). These results suggest that estrogen signalling regulates the intrinsic excitability in the embryonic brain through a long-term action of estrogen signalling rather than by a transient regulation of the neuronal activity. Next, we investigated whether EROB cells are involved in estrogen-mediated inhibition of excitability in the OB, and if so, whether ablation of EROB cells could prevent this estrogen mediated effect. To assess this, we used Tg(*ERE:nfsBmCherry*) x Tg(*elavl3/HuC:GCaMP6s*) embryos (details in “Methods”) that allows a simultaneous analysis of both EROB cell ablation and live GCaMP6s imaging in the same embryo (Fig. S3C). In EE2+MTZ exposed (EROB cell ablated) embryos, the level of GCaMP signals was restored to a level comparable with that observed in control embryos (Fig. 6D). Quantifications of the time-averaged GCaMP signals (detailed in “Methods”) revealed that EE2 exposure inhibited the intrinsic excitability both in specific glomerulus (mdGs and maG) and in the entire OB (Fig. 6Ei-iii, p* < 0.05 ANOVA with Tukey’s test). In contrast, EROB cell ablation by EE2+MTZ treatment prevented such EE2-induced neuronal inhibition (Fig. 6Ei-iii, p* < 0.05 ANOVA with Tukey’s test). These data confirmed that estrogen regulates the intrinsic neuronal activity via EROB cells.

**Figure 6.**
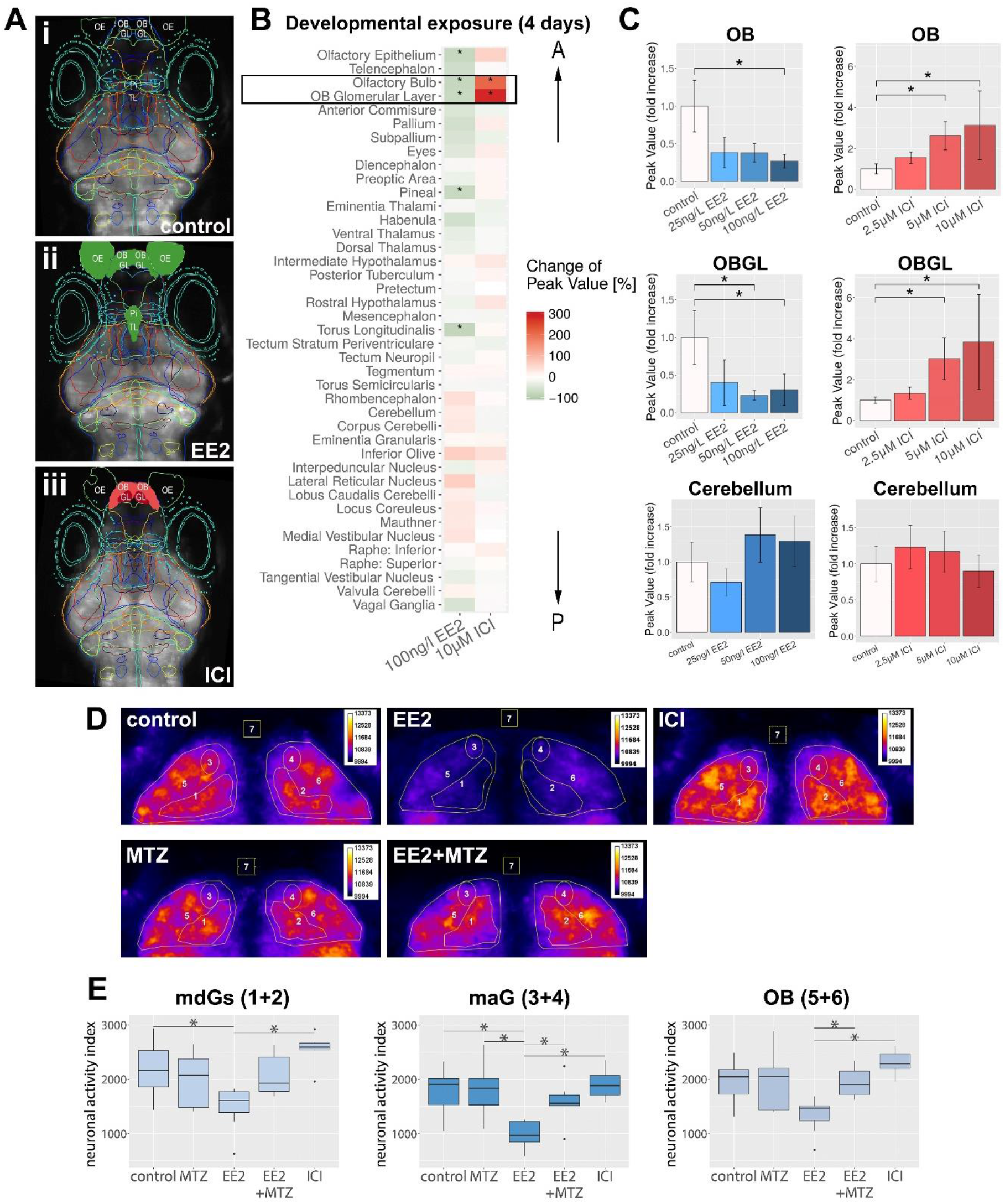
Estrogen/EROB cell cascade inhibits the intrinsic spontaneous neuronal activity in the olfactory bulb (OB). (A) LSM images of 4 dpf elavl3/HuC:GCaMP6s zebrafish embryo brain (dorsal view) of control (top), EE2 exposed (middle) and ICI exposed embryos (bottom) (see details in “Methods”). Each coloured line in the images represents a brain region of interest (ROI). ROIs whose GCaMP activity are inhibited by EE2 are coloured with green and those activated by ICI are coloured with red (based on the data in Fig.6B). Olfactory epithelia, OE; olfactory bulb, OB; olfactory bulb glomerular layer, OBGL; pineal, Pi and torus longitudinalis, TL. (B) Heat map displaying changes (%) of GCaMP signals in 41 different brain regions. A black rectangle indicates the most affected ROIs (OB and OBGL). The order of ROIs represents the location of a ROI along a-p axis (top-bottom). p* < 0.05, the Chi-square difference test with Tukey’s post-hoc test, N=8 each. (C) Dose-dependent effects of EE2 and ICI on intrinsic neuronal activity. p* < 0.05, Chi-square test with Tukey’s test, N=8 each. (D) Representative confocal z projection images (with Fire image mode) of time-averaged GCaMP6s signals in the OB of control, EE2, ICI (top row), MTZ, EE2+MTZ (bottom row) -treated 4 dpf Tg(*ERE:mCherry*) x Tg(*elavl3/HuC:GCaMP6s*) embryos. ROIs 1 and 2, left and right mdG1-6 (mdGs); ROIs 3 and 4, left and right maG; ROIs 5 and 6, left and right OB and ROI 7 for a background signal. (Ei-iii) Time-averaged mean intensities of GCaMP signals in mdGs (Ei: ROIs 1 and 2), in maG (Eii: ROIs 3 and 4) and in OB (Eiii: ROIs 5 and 6) from each experimental group (N=7-9) are shown. p* < 0.05, ANOVA with Tukey post-hoc test.

### Developmental exposure to EE2 or ICI abolishes olfactory-mediated avoidance responses

Finally, to examine the consequence of the change in the excitability of the OB induced by EE2 or ICI exposure, we next examined whether olfaction-mediated behaviour was affected by these exposures. By choosing to work with the fear-related infochemical cadaverine (Hussain et al., 2013), a death-associated odour, we aimed to determine if the disruption of the estrogen pathway during development interferes with the development of innate, olfactory avoidance that is crucial for survival (Fig. 7A and 7B). Using a video tracking system, we found no location preference for zebrafish larvae in either control, EE2 or ICI groups during the acclimation period (Fig. 7C, left column; Fig. 7D). After the administration of cadaverine, which is known to cause fish to swim away from the exposure zone (Hussain et al., 2013), avoidance behaviour was seen in the control group only (Fig. 7C, right column; Fig. 2E, p* < 0.05, Chi-square test with Tukey’s test). Conversely, neither EE2 or ICI-exposed larvae showed a clear location preference before or after cadavarine administration (Fig. 7C, 7D and 7E). In this system, we could not determine whether genetic ablation of EROB cells also alters olfactory-mediated avoidance response because MTZ treatment alone abrogated a cadavarine-mediated avoidance behaviour. Our data indicate that both activation (by EE2 exposure) and suppression (by ICI exposure) of estrogen signalling during brain development indeed affected the normal expression of fear-related behaviour triggered by sensing an alarm odour.

**Figure 7.**
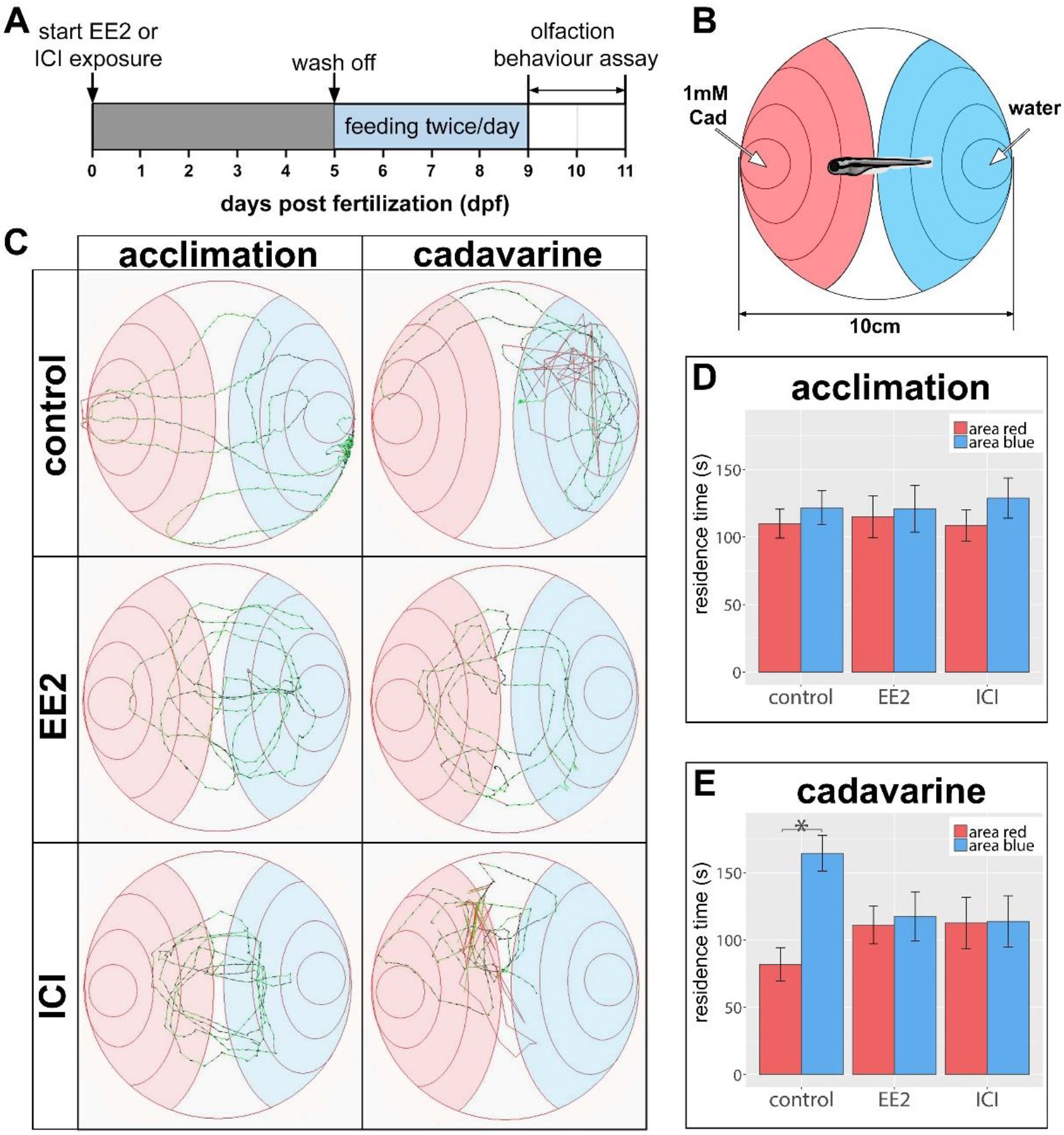
Developmental exposure to EE2 or ICI abolishes olfactory-mediated avoidance responses. (A) Schematic illustrating the protocol for the developmental exposure to EE2 or ICI and subsequent timing for the olfaction bioassay. (B) Illustration of an experimental chamber for the behaviour assay. Arrow indicates the administration sites of either cadavarine or water. Place preference of the larva was assessed by quantifying the residence time in the red (cadavarine +) or blue (water +) areas during the 5 minute assay period. (D) Representative swimming trajectories of control (top row), EE2 treated (middle row) and ICI treated (bottom row) larvae (details in “Methods”). Trajectories during acclimation (left column) and after cadavarine administration (right column) from the same larva are shown. (E and F). Averaged residence time in red (cadavarine) or blue (water) area before (E, acclimation) and after cadavarine administration (F, cadavarine) in control (n=14), EE2 (n=12) or ICI (n=12) treated larvae. p* < 0.05, Chi-square test with Tukey’s test.

## Discussion

### OB as a primary target of the estrogen signalling pathway in the zebrafish embryonic brain

Here we showed that the OB in the forebrain is a primary target of the estrogen signalling pathway in the zebrafish embryo. In our estrogen biosensor (ERE-TG) zebrafish lines, estrogen/ER-dependent transcriptional activation is shown as a reporter gene (e.g. GFP or mCherry) expression in a specific estrogen target cell/tissue, and such response is suppressed by the ER antagonist, ICI (Lee et al., 2012a). Using these ERE-TG lines, we found that a small number of glial cells in the OB, EROB cells, specifically respond to estrogen from shortly after the regional specification of the embryonic brain (i.e. 27 hpf) up to 10 dpf (Fig. 1). Interestingly, the ontogenic profiles of EROB cells coincide with the critical time window for olfactory glomerular development in the zebrafish embryo and larvae (Fig. 1J) (Braubach et al., 2013; Li et al., 2005; Miyasaka et al., 2013). This suggests that the estrogen/EROB cell cascade is required for olfactory glomerular development during early embryonic/larval stages. In contrast to the OB specific estrogen responsiveness seen here, the expression domains of ERs are not specific to the OB but rather occur more widely across the CNS in zebrafish embryos from 24 hpf onwards (Tingaud-Sequeira et al., 2004). Asking the question why did we detect such a localised estrogen activity in our ERE-TG lines given the widespread distribution of ERs from 24hpf?, we think the OB-specific estrogen activity may be attributed to the brain-region specific expression of *cyp19a1b* gene (the brain specific aromatase). We show that EROB cells express mCherry and GFP in the double transgenic embryo carrying both *ERE:mCherry* and *cyp19a1b:GFP* reporter genes (Fig. 2Bi-ii), indicating EROB cells express *cyp19a1b* gene. It is likely that *cyp19a1b* possesses a brain-region specific enhancer in conjunction with estrogen responsive element (ERE) in its transcriptional regulatory regions (Tong and Chung, 2003) and thereby could locally produce estrogen through an autocrine mechanism. The estrogen signalling cascade in EROB cells could therefore be amplified by a positive-feedback via their own aromatase activity. It is noteworthy that, unlike the *cyp19a1b:GFP* transgene which includes 3.4kb of the endogenous *cyp19a1b* promoter region (Tong et al., 2009), the *ERE:Gal4ff* transgene in our ERE-TG models contains three tandem-repeated short EREs combined with a TATA minimal promotor only. Thus, the ERE-TG models can detect ER-mediated transcriptional activities widely throughout the embryo without being affected by tissue specific enhancers or suppressors (Lee et al., 2012a). These newly identified EROB cells in ERE-TG models reveal possibly the most prominent estrogen activity commencing at the earliest stage of embryonic brain development, which is likely to be enhanced locally through the aromatase-mediated positive feedback mechanism.

We found that estrogen promotes the extension of EROB cell projections and facilitates the tight interaction between the EROB cells with OSN termini at OB glomeruli. Inhibiting the estrogen pathway by ICI markedly impairs the growth of EROB cell projections (Fig. 3). Both ICI and EROB cell ablation cause substantial defects in the olfactory glomerular development resulting in their reduced size, disruption of the pathfinding of OSN projections and altered topology of olfactory glomeruli in the OB (Fig. 4). Interestingly, previous work using immuno-histological analyses of rat embryonic brain section samples have suggested a role of embryonic radial glia cells (RGCs) in olfactory glomerular development (Bailey et al., 1999; Ramon-Cueto and Valverde, 1995; Valverde et al., 1992); RGCs occur along the midline (ventricle) in the telencephalon from an early embryonic stage (E11-14) and later (E17-18) project towards olfactory nerve fibre layer where OSN axons surround the majority of OB surface. Subsequently (E19-20), the termini of RGCs intermingle with OSN axons to form the foundation of OB glomeruli, protoglomeruli, in the olfactory glomerular layer (Bailey et al., 1999; Ramon-Cueto and Valverde, 1995; Valverde et al., 1992). These observations support a conserved developmental mechanism for the estrogen/EROB cell cascade between fish and mammals. In fact, EROB cells may be a type of such embryonic RGCs given that a brain specific aromatase (*cyp19a1b*) is induced in EROB cells in response to estrogen and this is known to be expressed specifically in a subpopulation of RGCs in zebrafish (Brion et al., 2012; Menuet et al., 2005). We found that suppression of estrogen signalling causes developmental abnormalities in key cellular and functional components of the olfactory glomerular layer, including the EROB cells, OSNs and inhibitory synapses: In ICI treated embryos, EROB cell projection networks are not established or maintained (Fig. 3). Here we show that without the EROB projection networks, which likely to be a scaffold of the olfactory glomerular layer, OSN axonal extension to OB glomeruli is markedly altered. Illustrating this, in ICI treated embryos, OSN termini are often ectopically positioned or fail to reach to the expected glomerulus (Fig.4). Similar phenotypes were also seen in EROB cell-ablated embryos (Fig.4). The defects in organisation of OB glomeruli can consequently impair the establishment of inhibitory synapses in OB glomeruli, which we were able to show for ICI treated embryos (Fig. 5).

### Estrogen and EROB cells establish the responsiveness of the embryonic olfactory system

We found that the OB specific roles of EE2/EROB cells in olfactory glomerular development, including the role in the establishment of olfactory inhibitory synapses, contribute also to neurophysiological processes: A developmental exposure of the zebrafish embryos with EE2 inhibits the intrinsic excitability predominantly in the OB whereas ICI activates the intrinsic excitability exclusively in the OB. The intrinsic neuronal activities in other brain regions are not affected by EE2 or ICI (Fig. 6). It is important to note that previous work has shown that in zebrafish embryos, elavl3/HuC promoter induces a fluorescent reporter protein predominantly in mitral cells, which is an excitatory output neuron in the OB (Li et al., 2005). We thus think that GCaMP signals in the OB we observe here most likely originate from the excitatory input in the OB, not the inhibitory input from interneurons.

Olfaction is essential for survival, for behaviours such as feeding, mating, social interaction and assessing dangers. Zebrafish embryos hatch after 48 hpf and start feeding at 4-5 dpf and odour-induced neuronal activation in the OB can be detected from as early as 2.5-3 dpf (Li et al., 2005). Olfaction-mediated behavioural response are observed from 4-5 dpf ((Lindsay and Vogt, 2004) and Takesono A. in preparation). In olfactory sensory systems, the intrinsic neuronal activity in the OB is known to be the basal excitability which contributes to the signal-to-noise ratio between the spontaneous and an odour (i.e. cadavarine)-evoked neuronal activities (Savigner et al., 2009). Thus an appropriate tuning of the intrinsic spontaneous excitability in the OB may be essential for odour information processing and for the subsequent behaviour. In fact, we found that a considerably lowered (by EE2) or excessively heightened (by ICI) intrinsic neuronal activity results in the loss of odour-induced behavioural response (Fig. 7). The intrinsic spontaneous excitability in the OB is also known to be required for precise targeting of OSN axons to appropriate olfactory glomeruli during olfactory glomerular development (Yu et al., 2004) as well as for refining the dendrite connectivity of a mitral cell (olfactory output neurons) to a specific olfactory glomerulus (Fujimoto, 2019). Thus, it may be an interesting perspective that estrogen-mediated modulation of the intrinsic neuronal activity in the OB could also affect the development of olfactory glomeruli, and vice versa.

Based on these results, we propose that estrogen/EROB cascade plays a key instructive role in the development of olfactory glomeruli in the OB: Estrogen primarily activates the growth of the EROB cell projection networks in the OB. EROB cells in turn act as a scaffold to guild the extension of OSN projections into olfactory glomeruli and facilitate the establishment of olfactory glomeruli including the local inhibitory circuits. Consequently, an estrogen/ERO cell cascade controls the intrinsic spontaneous excitability in the OB that determines the olfaction-mediated behaviour.

### Estrogen/EROB cascade – a potential site of action for EDCs or a possible link to neurodevelopmental disorders

A wealth of data exist on the effect of exposure on olfaction-mediated responses, including on odour-induced neurogenesis, excitability and olfaction-linked cognition/memory and behaviour in mammals including human (Brus et al., 2016; Cherian et al., 2014; Doty et al., 2008; Kollndorfer et al., 2016; Veyrac and Bakker, 2011). These studies, however, have primarily focused on adult stages of human and rodents with very little attention given to the role of estrogen in development of olfactory sensory system at embryonic stages. Our data further highlights the possibility that the estrogen/EROB cell cascade in early embryonic stages could be an important site of action for EDCs in environment that interfere with the endogenous functions of estrogen. If so, exposure to environmental estrogens during embryonic brain development could cause adverse effects on olfactory sensory system and impact on fitness and social/sexual behaviour in later life. In fish, chemical communication through olfaction also plays a crucial role in shaping ecological interactions, allowing animals to locate food sources, predators, habitats or mates (Wyatt, 2003). In fact, we have very recently found that EROB cells respond also to the estrogenic endocrine disruptors, nonylphenol, bisphenol A and genestin, resulting in very similar ontogenic profiles as reported here for exposure to EE2 (A.Takesono et al., in preparation). Given that the developing embryo is generally the most susceptible life stage to environmental toxicants, this possibility warrants further investigation.

Furthermore, alterations in estrogen/EROB cascade during development might link with certain neurodevelopmental disorders (i.e. ADHD and autism) which have been implicated with an imbalance in sex hormone-mediated signalling pathways (Crider and Pillai, 2017; Mustieles et al., 2015). Notably, the clinical symptoms for these conditions are often associated with olfactory impairment (anosmia) and/or altered olfactory sensitivity (Endevelt-Shapira et al., 2018; Sundermann et al., 2008; Valdes-Socin et al., 2014). Dysfunctions in estrogen cascade could therefore be related to the pathogenesis of such neurodevelopmental disorders. Interestingly, estrogenic compounds have been shown to selectively rescue the behaviour phenotype observed in *Contactin Associated Protein-like 2* (*CNTNAP2*) mutant zebrafish embryo, an autism-related gene mutant line carrying GABAergic neuron deficits (Hoffman et al., 2016). Thus, it would be interesting to examine whether autism phenotypes in CNTNAP2 mutant could relate to estrogen/EROB cell cascade.

In summary, our work identifies a fundamental role of estrogen in development of olfactory sensory system in embryonic brain, which could result in a long-lasting influence on neuronal circuitries responsible for olfaction and could impact on behaviour and fitness in later life. Further research is required to determine if this estrogen/EROB cell cascade is conserved across vertebrate species. The identification of specific target genes involved in estrogen/EROB cell cascade would be the next step to elucidate a precise mechanism of action of estrogen in EROB cells.

## Acknowledgments

This work was funded by Biotechnology and Biological Sciences Research Council (BBSRC) project grant BB/L020637/1 and BBSRC Japan Partnering Award BB/P025528/1. P.S. was funded by Erasmus Internship at UEXE. We thank Olivier Kah for providing the Tg(*cyp19a1b:GFP*), his support in training AT in immunohistochemistry and for his valuable discussions, Izumi Fukunaga for her generous help for co-localisation analysis and for critical comments on the manuscript and Robert Heathcote for his advice on analysing the behaviour data. We also thank Htoo Wai for his technical help in the initial work establishing transgenic lines, Nobuhiko Miyasaka for providing the *pT2A UAS:DsRed-Ex* plasmid DNA. We are grateful to ARC staff at University of Exeter for technical support in the husbandry of the zebrafish and Gasper Jekely and Steffen Schlopp for critical reading of our manuscript.

## Author contributions

Conceptualization, A.T., T.K. and C.R.T.; Methodology, A.T., P.S., M.J.W., T.K. and C.R.T; Formal analysis, P.S.; Investigation, A.T., P.S.; Writing – Original Draft, A.T; Writing – Review & Editing, A.T., P.S., M.J.W., T.K. and C.R.T.; Funding Acquisition, A.T., T.K. and C.R.T.; Resources, A.T., A.S., J.M.G, O.L., T.K and C.R.T.; Supervision, A.T., T.K. and C.R.T.

## Corresponding author

Correspond to Charles R. Tyler and Aya Takesono

## Methods

Fish husbandry, TG zebrafish lines used, immunohistochemistry/imaging analyses and all other methods not detailed here are described in Supplemental Methods.

### Chemical exposures

#### Developmental exposure to estrogen or an estrogen receptor antagonist (ICI)

To examine the roles of estrogen signalling pathway in brain development, zebrafish embryos collected from the selected transgenic zebrafish lines, depending on the requirement for the different experiments, were exposed to, 100 ng/L ethinylestradiol (EE2) or 10μM ICI 182,780 (ICI, an estrogen receptor antagonist, Sigma-Aldrich)(aqueous exposure in system water) from 1 to 96 hours post fertilisation (hpf). At 96 hpf, chemically exposed embryos were washed once and used for immunohistochemistry (Fig. 4 and 5) and GCaMP imaging (Fig.6). For the dose response experiments to these compounds (also Fig.6) zebrafish embryos were exposed to 25, 50 or 100 ng/L EE2 or to 2.5, 5 or 10 μM ICI. For the olfaction behaviour assay, the exposure period to these chemicals was extended to 5 days (up to 120hpf).

#### Transient exposure to estrogen or estrogen receptor antagonist

For testing the acute effect of exposure to estrogen (EE2) or the estrogen receptor antagonist (ICI) on neuronal activity (Fig. S5), elavl3/HuC:GCaMP6s embryos were cultured in aquarium system water without chemical exposure up to 96 hpf. At 96 hpf, prior to imaging with light sheet microscopy (LSM), embryos were exposed to 100 ng/L EE2 or 10 μM ICI in 1 x E3 media for 20 minutes at room temperature. Following exposure, GCaMP6s signals in each embryo were examined by LSM, as described below.

#### Chemical exposure conditions for chemical/genetic ablation of EROB cells in ERE:mCherry line

To induce selective ablation of EROB cells, we employed nitroreductase-mediated chemical/genetic cell ablation (Curado et al., 2008). In the ERE:mCherry line, Tg(*ERE:Gal4ff; UAS:nfsBmCherry*), intracellular nitroreductase synthesis is under control of estrogen-mediated ERE-activation. Selective ablation of EROB cells was achieved by applying EE2 together with a prodrug metronidazole (MTZ) to induce a cytotoxic metabolites only in estrogen responsive cells. ERE: *nfsB*mCherry embryos were exposed to either aquarium system water (control), 100 ng/L EE2, 7.5 mM MTZ (Sigma-Aldrich) or 100 ng/L EE2 plus 7.5 mM MTZ (EE2+MTZ) from 1 to 33 hpf. This mixture treatment condition of EE2 in combination with MTZ was established as the optimised condition for effective ablation of EROB cells in the absence of developmental abnormalities (50% reduction of EROB cells at 4 dpf, Fig. S4A). Following the chemical exposures, embryos at 33hpf were washed once and cultured in aquarium water until 4 dpf and subsequently used for immunohistochemistry (Fig.4) and GCaMP6s imaging (Fig.6).

### EROB cell projection analyses

To examine the effect of estrogen on cellular projections of the EROB cells, ERE:GFP embryos were treated with 100ng/L EE2 from 1 hpf to 72 hpf to induce GFP in EROB cells and they then underwent either a washout period from 72hpf to 120hpf (in aquarium water, referred as “control”) or a washout period in aquarium water that also included 10μM ICI to inhibit estrogen activity over the same time period (72hpf to 120hpf, referred to as “ICI”). At 120hpf, chemical exposed embryos were washed once and then mounted in 0.7% low melting point agarose in a 35 mm diameter glass-bottom dish (MatTek) with the embryo angled at approximately 40° dorsal surface facing downwards. Confocal images of EROB cells were obtained using Zeiss 880 Airyscan panel acquisition with x40 objective. Images were airy-processed and stitched using Zeiss Zen Black software and further processed using Fiji with a set parameters for brightness/Contrast adjustment and background subtraction. These images were then presented as max intensity projection images. To quantify the effect of ICI on the EROB cell projection networks, mean grey values of GFP signals within the margins of the EROB cells (as marked with a green square in Fig. 3Ai or 3Bi) were established using surface plot function in Fiji. Mean GFP signals within the EROB cell margin were also plotted along the width of the OB margin, unifying individual measurements by assigning the position of the mid-line to 0 and the ends of the OB pia to 1 (right) and −1 (left). The average of grey values and sum of grey values (% of total grey value in each embryo) in both edges of the EROB cell margin ([−1:−0.7], left margin; [0.7:1], right margin) were analysed to examine the role of estrogen activity for establishment of EROB cell projections (N=6 for each).

#### Statistics

Statistics were performed with R (version 3.6.1) (Team, 2019). A Shapiro-Wilk test identified the data for the average of grey values in both left and right margins were not normally distributed and applying Levene’s test showed heterogeneity of variances. Thus, a non-parametric Mann Whitney U test was performed. The same statistical analysis was also applied for the fraction of sum of grey value in both margins. The EROB cell count showed normal distribution and homogeneity of variances, thus a t-test was performed to explore the effect of ICI on the number of EROB cells.

### Immunohistochemical analysis for Gephyrin expressing Inhibitory synapses

Chemical exposure conditions for ERE:GFP embryos were as described above. At 4dpf, exposed embryos were anaesthetised with 0.03% NM222 and then embedded in NEG-50 solution (ThermoFisher Scientific) without PFA fixation, flash frozen in liquid nitrogen and stored at −80°C. Embedded frozen samples were transverse-sectioned in consecutive 25μm thickness sections using a CM1950 cryostat (Leica). Sections were transferred to Fisher Plus Gold slides and post-fixed with 4% PFA/PBS for 15min at room temperature. Fixed sections were gently rinsed two times with 1xPBS and treated with pepsin reagent (Sigma-aldrich, R2283) for 5 min at 37°C, followed by carefully rinsing two times with 1xPBS. Subsequently, the samples were permeabilised with 0.1% Triton/PBS for 3 min at room temperature and then rinsed twice with 1xPBS and blocked with 2% BSA/0.2% milk in PBS for one hour at room temperature. Blocked sections were subsequently stained with mouse anti-gephyrin antibody (1:500 in the same blocking solution, synaptic systems, #147011) and rabbit anti-GFP antibody (1:500 in blocking, AMS Biotechonology) at 4°C overnight. The samples were then washed three times with 1xPBS and incubated with Alexa 594 conjugated anti-mouse IgG antibody (1:500 in blocking buffer, ThermoFisher Scientific, A11012) and Alexa 488 conjugated anti-rabbit IgG antibody (1:500 in blocking buffer, ThermoFisher Scientific, A11034) for one hour at room temperature. The stained samples were then washed three times with 1xPBS and incubated with Hochest 33342 (1:25,000 in 1 x PBS, ThermoFisher Scientific) for 30 minutes. Finally, the samples were washed twice with 1 x PBS and mounted with ProLong Gold antifade reagent (ThermoFisher scientific)._Stained sections were imaged using Airy scan Zeiss LSM880 with x20 or x40 objective lens. Th region covering the whole olfactory bulb was captured from 3 or 4 consecutive (25μm thickness) sections per individual sample. Images were airy-processed and/or stitched (for panel acquisition images) using Zeiss Zen black software and further processed with Fiji with a set parameters for brightness/Contrast adjustment and background subtraction. Gephyrin puncta were quantified by applying a threshold (IJ_IsoData method) and watershed algorithm to obtain binary images of gephrin + puncta and selecting only those ranging in the sizes 0.1-0.4, 0.41-0.99 and 1-10 μm^2^ using the analyze particles function in Fiji. The outlined drawings of the size selected puncta were merged with the nuclei staining image of the section to verify the positions of the puncta in the OB regions. Positions of the OB in the brain sections were confirmed in accordance with Atlas of early zebrafish brain development (Wullimann, 2015). The number of puncta in the OB regions was counted manually using multipoint function in Fiji. Gephyrin puncta numbers in each puncta-size range group were summed and displayed as % of total number in control (N=4 each).

#### Statistics for gephyrin puncta analysis

Statistics were performed with R (version 3.6.1)(Team, 2019). Normality of the data was confirmed with the Shapiro test. Levene’s test showed homogeneity of variances and a linear model was built. A one-way ANOVA, in conjunction with Tukey’s post-hoc test, was performed for pair-wise comparisons of the treatments, using the ‘multcomp’ package in R.

### GCaMP6s imaging

#### Experimental settings for GCaMP6s imaging using Light Sheet Microscopy (LSM)

The detailed procedures for sample preparation for GCaMP6s imaging and for imaging acquisition using a custom-built LSM are described in (Winter et al., 2017). A full brain volume of GCaMP6s images for zebrafish embryo larvae is captured in around 1.8 secs (10 horizontal plane optical z-slices in 24 μm steps). These were taken repeatedly for around 6 mins from which the region of interest (ROI)-specific neuronal activity data was extracted. 41 anatomically registered brain ROIs were included which encompassed all the major brain structures of relevance. “Peak value” of GCaMP6s signals in a ROI were compared statistically between control and experimental groups (N=8 for each). “Peak value” represents the average of the peaks of GCaMP6s signals within that ROI over the experimental period and are considered the most revealing parameter for showing changes in the intrinsic neuronal activity within each brain region (Winter et al., 2017). To minimise experimental subject variability in GCaMP6s fluorescence, elavl3/HuC:GCaMP6s embryos were pre-screened for a similar basal GCaMP expression level in the brain before LSM imaging. All experimental treatments were conducted within one batch of embryos, and repeated on two separate occasions, to account for possible batch to batch variations and all imaging conditions were kept identical throughout the study (Winter et al., 2017).

#### Statistics for LSM GCaMP data

Statistics were performed with R (version 3.2.3-4). The LSM data, after processing through the Python image pipeline, was analysed using a generalised linear model with ‘Treatment’, ‘Region of interest (ROI)’ and the interaction of both as fixed effects. A Shapiro-Wilk test showed the data were not normally distributed (p*< 10^-16^) and the box cox normality plot suggested a logarithmic correlation. Therefore, a gamma distribution with a logarithmic link was used to fit the continuous data. The Chi-square difference test in conjunction with Tukey’s post-hoc test were performed using the estimated marginal means ‘emmeans’ package in R.

#### GCaMP imaging using Zeiss airyscan

To analyse neuronal activity under the condition of EROB cell ablation, heterozygous embryos of Tg(*ERE:Gal4ff; UAS:nfsBmCherry*) x Tg(*elavl3/HuC:GCaMP6s*) were produced from a pair-cross of homozygous parent of each TG fish. Chemical exposure conditions for EROB cell ablation and for inhibiting ER activation by ICI (10μM) are described above. An efficient ablation of EROB cells were confirmed as shown in Fig. S4B. We emphasise that this system had a reduced (50%) sensitivity for GCaMP6s imaging, as compared with that of homozygous of Tg(*elavl3/HuC:GCaMP6s*), due to the heterozygosity of the transgenes. To minimise the variation, all datasets were acquired on the same date using the same batch of embryos (n=7 in each experimental group). The same experiments were repeated twice with similar results. At 96 hpf, exposed embryos were washed once with 1 x E3. Each embryo was treated with 4 mM anti-nicotinic neuromuscular blocker tubocurarine (4mM)(Sigma-Aldrich) until muscle tone was lost. The immobilised embryo was then quickly mounted in 0.7% low melting point agarose in a 35 mm diameter glass-bottom dish (MatTek) with the embryo angled at approximately 40° with the dorsal surface facing downwards. Live imaging of the elavl3/HuC:GCaMP6s was carried out using Zeiss 880 in fast acquisition mode with Airyscan, which achieves 9 consecutive optical z section images extending through the entire OB (scan depth 63 μm, 7 μm step each) in 2.2 sec, allowing active neurons labelled with elavl3/HuC:GCaMP6s throughout glomerular layer to be detected. This approach allowed individual cell-level functional imaging to be undertaken at a similar temporal resolution to that obtained using LSM, but for a lower z-depth. This was nonetheless appropriate for imaging the olfactory bulb region, rather than the whole brain. Image acquisition was repeated for 50 cycles, generating 50 × 9 optical z section images. Z projection images of EROB cells (ERE:mCherry) were also obtained from the same experimental embryos to confirm nitroreductase-mediated EROB cell ablation.

#### Data processing of airyscan GCaMP6s images and statistics

Raw data for elavl3/HuC:GCaMP6s images from each experimental embryo were processed in Fiji to select the z projections from period between 15 to 35 cycles and to obtain average intensity z projection images, and then analysed for a region specific fluorescence (GCaMP6s) intensity using the region of interest (ROI) manager. ROIs for the medio-anterior glomerulus (maG), the medio-dorsal glomeruli (mdG1-6), as well as the entire OB, were manually selected using the polygon selection tool. For each ROI, fluorescence intensity (FI) data were presented as the time-averaged mean FI. The statistics were performed with R (version 3.2.3-4). Normality of the data was confirmed with the Shapiro-Wilk test and Levene’s test showed homogeneity of variance. Hence, a linear model was built. The one-way ANOVA in conjunction with Tukey’s post-hoc test was performed for pair-wise comparisons of the treatments, using the ‘multcomp’ package in R.

### Olfactory behaviour assay

Chemical exposure to ERE:mCherry embryo were performed as described above. At 5dpf, EE2 or ICI exposed embryos were washed and cultured in a glass dish containing 75 ml of aquarium system water at a density of 30 embryo-larvae/75 ml water and were then fed until the day of the olfaction behaviour assay at 9-11 dpf (Fig. 7A). One third of the volume of incubation water was changed daily until the time of the behaviour assessment. Larval movement was measured using automated videotracking (ViewPoint, France (Readman et al., 2013; Winter et al., 2008)) in an experimental arena (Fig. S2B). Each larva was assessed prior to testing for olfactory responsiveness to ensure an appropriate level of spontaneous swimming behaviour. This was achieved by placing each larva into the experimental arena, which contained 45ml of fresh aquarium water, and recording its swimming behaviour for a 5 minute acclimation period at 25 frames per second (fps). Individuals entering more than 7 areas without a location bias or more than 6 areas with a location bias to the centre of the chamber (the location areas in the experimental chamber are shown in Fig. 7B) were considered appropriate for the olfactory test. Subsequently, 200μl of 1mM cadavarine was introduced gently through a silicon capillary tube into one side of the experimental arena, while the same amount of water was simultaneously applied to the opposite side of the chamber. This produced a cross-arena biased gradient of cadavarine with minimum disturbance of the animal. During method development, it was established that the gradient of a chemical was maintained throughout the 5 minutes of recording based on the diffusion rate of phenol red (data now shown). The swimming behaviour of the larva after cadavarine administration was recorded for a further 5 minutes and the data analysed for the time spent within the cadaverine-treated or untreated areas using Viewpoint software. The cadavarine-mediated avoidance response was defined as the average duration spent (residence time) in the red (cadavarine high) versus blue (cadavarine low) areas (Fig. 7E).

#### Statistics for behaviour data

Statistics were performed with R (version 3.2.3-4). The paired experimental design and thus dependency of the test parameters on the individuality of each zebrafish larvae required a mixed effect model analysis approach using the ‘individual’ as a random effect. The distribution of the data was best described with a normal distribution, leading to the choice of a linear mixed effect model, using the ‘lme4’ package in R. Fixed effects of this model were Treatment and Area as well as their two-way interaction. The date of the experiment and the developmental stage (9-11 days) had no significant effect on the model. For pair-wise comparisons within the treatment groups, Tukey’s test was performed using the ‘emmeans’ package in R.

## References

Bailey, M. S., Puche, A. C. and Shipley, M. T. (1999). Development of the olfactory bulb: evidence for glia-neuron interactions in glomerular formation. J Comp Neurol 415, 423–448.

Batista-Brito, R., Close, J., Machold, R. and Fishell, G. (2008). The distinct temporal origins of olfactory bulb interneuron subtypes. J Neurosci 28, 3966–3975.

Bondesson, M., Hao, R., Lin, C. Y., Williams, C. and Gustafsson, J. A. (2015). Estrogen receptor signaling during vertebrate development. Biochim Biophys Acta 1849, 142–151.

Braubach, O. R., Miyasaka, N., Koide, T., Yoshihara, Y., Croll, R. P. and Fine, A. (2013). Experience-dependent versus experience-independent postembryonic development of distinct groups of zebrafish olfactory glomeruli. J Neurosci 33, 6905–6916.

Braun, J. M., Yolton, K., Dietrich, K. N., Hornung, R., Ye, X., Calafat, A. M. and Lanphear, B. P. (2009). Prenatal bisphenol A exposure and early childhood behavior. Environ Health Perspect 117, 1945–1952.

Brion, F., Le Page, Y., Piccini, B., Cardoso, O., Tong, S. K., Chung, B. C. and Kah, O. (2012). Screening estrogenic activities of chemicals or mixtures in vivo using transgenic (cyp19a1b-GFP) zebrafish embryos. PLoS One 7, e36069.

Brus, M., Trouillet, A. C., Hellier, V. and Bakker, J. (2016). Estradiol-induced neurogenesis in the female accessory olfactory bulb is required for the learning of the male odor. J Neurochem 138, 457–468.

Cherian, S., Wai Lam, Y., McDaniels, I., Struziak, M. and Delay, R. J. (2014). Estradiol rapidly modulates odor responses in mouse vomeronasal sensory neurons. Neuroscience 269, 43–58.

Crider, A. and Pillai, A. (2017). Estrogen Signaling as a Therapeutic Target in Neurodevelopmental Disorders. J Pharmacol Exp Ther 360, 48–58.

Curado, S., Stainier, D. Y. and Anderson, R. M. (2008). Nitroreductase-mediated cell/tissue ablation in zebrafish: a spatially and temporally controlled ablation method with applications in developmental and regeneration studies. Nat Protoc 3, 948–954.

Della Torre, S., Rando, G., Meda, C., Ciana, P., Ottobrini, L. and Maggi, A. (2018). Transcriptional activity of oestrogen receptors in the course of embryo development. J Endocrinol 238, 165–176.

Derouiche, L., Keller, M., Martini, M., Duittoz, A. H. and Pillon, D. (2015). Developmental Exposure to Ethinylestradiol Affects Reproductive Physiology, the GnRH Neuroendocrine Network and Behaviors in Female Mouse. Front Neurosci 9, 463.

Doty, R. L., Kisat, M. and Tourbier, I. (2008). Estrogen replacement therapy induces functional asymmetry on an odor memory/discrimination test. Brain Res 1214, 35–39.

Endevelt-Shapira, Y., Perl, O., Ravia, A., Amir, D., Eisen, A., Bezalel, V., Rozenkrantz, L., Mishor, E., Pinchover, L., Soroka, T., et al. (2018). Altered responses to social chemosignals in autism spectrum disorder. Nat Neurosci 21, 111–119.

Fan, X., Warner, M. and Gustafsson, J. A. (2006). Estrogen receptor beta expression in the embryonic brain regulates development of calretinin-immunoreactive GABAergic interneurons. Proc Natl Acad Sci U S A 103, 19338–19343.

Fujimoto, S., Marcus N. Leiwe, Richi Sakaguchi, Yuko Muroyama, Reiko Kobayakawa, Ko Kobayakawa, Tetsuichiro Saito, Takeshi Imai (2019). Spontaneous activity generated within the olfactory bulb establishes the discrete wiring of mitral cell dendrites. bioRxiv.

George, F. W. and Ojeda, S. R. (1982). Changes in aromatase activity in the rat brain during embryonic, neonatal, and infantile development. Endocrinology 111, 522–529.

Gioiosa, L., Parmigiani, S., Vom Saal, F. S. and Palanza, P. (2013). The effects of bisphenol A on emotional behavior depend upon the timing of exposure, age and gender in mice. Horm Behav 63, 598–605.

Godfray, H. C. J., Stephens, A. E. A., Jepson, P. D., Jobling, S., Johnson, A. C., Matthiessen, P., Sumpter, J. P., Tyler, C. R. and McLean, A. R. (2019). A restatement of the natural science evidence base on the effects of endocrine disrupting chemicals on wildlife. Proc Biol Sci 286, 20182416.

Green, J. M., Metz, J., Lee, O., Trznadel, M., Takesono, A., Brown, A. R., Owen, S. F., Kudoh, T. and Tyler, C. R. (2016). High-Content and Semi-Automated Quantification of Responses to Estrogenic Chemicals Using a Novel Translucent Transgenic Zebrafish. Environ Sci Technol 50, 6536–6545.

Hoffman, E. J., Turner, K. J., Fernandez, J. M., Cifuentes, D., Ghosh, M., Ijaz, S., Jain, R. A., Kubo, F., Bill, B. R., Baier, H., et al. (2016). Estrogens Suppress a Behavioral Phenotype in Zebrafish Mutants of the Autism Risk Gene, CNTNAP2. Neuron 89, 725–733.

Hussain, A., Saraiva, L. R., Ferrero, D. M., Ahuja, G., Krishna, V. S., Liberles, S. D. and Korsching, S. I. (2013). High-affinity olfactory receptor for the death-associated odor cadaverine. Proc Natl Acad Sci U S A 110, 19579–19584.

Kawai, K., Nozaki, T., Nishikata, H., Aou, S., Takii, M. and Kubo, C. (2003). Aggressive behavior and serum testosterone concentration during the maturation process of male mice: the effects of fetal exposure to bisphenol A. Environ Health Perspect 111, 175–178.

Kinch, C. D., Ibhazehiebo, K., Jeong, J. H., Habibi, H. R. and Kurrasch, D. M. (2015). Low-dose exposure to bisphenol A and replacement bisphenol S induces precocious hypothalamic neurogenesis in embryonic zebrafish. Proc Natl Acad Sci U S A 112, 1475–1480.

Kollndorfer, K., Ohrenberger, I. and Schopf, V. (2016). Contraceptive Use Affects Overall Olfactory Performance: Investigation of Estradiol Dosage and Duration of Intake. PLoS One 11, e0167520.

Konkle, A. T. and McCarthy, M. M. (2011). Developmental time course of estradiol, testosterone, and dihydrotestosterone levels in discrete regions of male and female rat brain. Endocrinology 152, 223–235.

Kramar, E. A., Chen, L. Y., Brandon, N. J., Rex, C. S., Liu, F., Gall, C. M. and Lynch, G. (2009). Cytoskeletal changes underlie estrogen’s acute effects on synaptic transmission and plasticity. J Neurosci 29, 12982–12993.

Lau, E. S.-W., Zhang, Z., Qin, M. and Ge, W. (2016). Knockout of Zebrafish Ovarian Aromatase Gene (cyp19a1a) by TALEN and CRISPR/Cas9 Leads to All-male Offspring Due to Failed Ovarian Differentiation. Scientific Reports 6, 37357.

Lee, O., Takesono, A., Tada, M., Tyler, C. R. and Kudoh, T. (2012a). Biosensor zebrafish provide new insights into potential health effects of environmental estrogens. Environ Health Perspect 120, 990–996.

Lee, O., Tyler, C. R. and Kudoh, T. (2012b). Development of a transient expression assay for detecting environmental oestrogens in zebrafish and medaka embryos. BMC Biotechnol 12, 32.

Li, J., Mack, J. A., Souren, M., Yaksi, E., Higashijima, S., Mione, M., Fetcho, J. R. and Friedrich, R. W. (2005). Early development of functional spatial maps in the zebrafish olfactory bulb. J Neurosci 25, 5784–5795.

Lichtensteiger, W., Bassetti-Gaille, C., Faass, O., Axelstad, M., Boberg, J., Christiansen, S., Rehrauer, H., Georgijevic, J. K., Hass, U., Kortenkamp, A., et al. (2015). Differential gene expression patterns in developing sexually dimorphic rat brain regions exposed to antiandrogenic, estrogenic, or complex endocrine disruptor mixtures: glutamatergic synapses as target. Endocrinology 156, 1477–1493.

Lindsay, S. M. and Vogt, R. G. (2004). Behavioral responses of newly hatched zebrafish (Danio rerio) to amino acid chemostimulants. Chem Senses 29, 93–100.

Liu, F., Day, M., Muniz, L. C., Bitran, D., Arias, R., Revilla-Sanchez, R., Grauer, S., Zhang, G., Kelley, C., Pulito, V., et al. (2008). Activation of estrogen receptor-beta regulates hippocampal synaptic plasticity and improves memory. Nat Neurosci 11, 334–343.

McCarthy, M. M. (2008). Estradiol and the developing brain. Physiol Rev 88, 91–124.

Menuet, A., Pellegrini, E., Brion, F., Gueguen, M. M., Anglade, I., Pakdel, F. and Kah, O. (2005). Expression and estrogen-dependent regulation of the zebrafish brain aromatase gene. J Comp Neurol 485, 304–320.

Miyasaka, N., Wanner, A. A., Li, J., Mack-Bucher, J., Genoud, C., Yoshihara, Y. and Friedrich, R. W. (2013). Functional development of the olfactory system in zebrafish. Mech Dev 130, 336–346.

Mukherjee, J., Cardarelli, R. A., Cantaut-Belarif, Y., Deeb, T. Z., Srivastava, D. P., Tyagarajan, S. K., Pangalos, M. N., Triller, A., Maguire, J., Brandon, N. J., et al. (2017). Estradiol modulates the efficacy of synaptic inhibition by decreasing the dwell time of GABAA receptors at inhibitory synapses. Proc Natl Acad Sci U S A 114, 11763–11768.

Mustieles, V., Perez-Lobato, R., Olea, N. and Fernandez, M. F. (2015). Bisphenol A: Human exposure and neurobehavior. Neurotoxicology 49, 174–184.

Nagayama, S., Homma, R. and Imamura, F. (2014). Neuronal organization of olfactory bulb circuits. Front Neural Circuits 8, 98.

Nash, J. P., Kime, D. E., Van der Ven, L. T., Wester, P. W., Brion, F., Maack, G., Stahlschmidt-Allner, P. and Tyler, C. R. (2004). Long-term exposure to environmental concentrations of the pharmaceutical ethynylestradiol causes reproductive failure in fish. Environ Health Perspect 112, 1725–1733.

Ogawa, S., Eng, V., Taylor, J., Lubahn, D. B., Korach, K. S. and Pfaff, D. W. (1998a). Roles of estrogen receptor-alpha gene expression in reproduction-related behaviors in female mice. Endocrinology 139, 5070–5081.

Ogawa, S., Washburn, T. F., Taylor, J., Lubahn, D. B., Korach, K. S. and Pfaff, D. W. (1998b). Modifications of testosterone-dependent behaviors by estrogen receptor-alpha gene disruption in male mice. Endocrinology 139, 5058–5069.

Porseryd, T., Volkova, K., Reyhanian Caspillo, N., Källman, T., Dinnetz, P. and Porsh Hällström, I. (2017). Persistent Effects of Developmental Exposure to 17α-Ethinylestradiol on the Zebrafish (Danio rerio) Brain Transcriptome and Behavior. Front Behav Neurosci 11, 69.

Ramon-Cueto, A. and Valverde, F. (1995). Olfactory bulb ensheathing glia: a unique cell type with axonal growth-promoting properties. Glia 14, 163–173.

Readman, G. D., Owen, S. F., Murrell, J. C. and Knowles, T. G. (2013). Do fish perceive anaesthetics as aversive? PLoS One 8, e73773.

Rissman, E. F., Heck, A. L., Leonard, J. E., Shupnik, M. A. and Gustafsson, J. A. (2002). Disruption of estrogen receptor beta gene impairs spatial learning in female mice. Proc Natl Acad Sci U S A 99, 3996–4001.

Savigner, A., Duchamp-Viret, P., Grosmaitre, X., Chaput, M., Garcia, S., Ma, M. and Palouzier-Paulignan, B. (2009). Modulation of spontaneous and odorant-evoked activity of rat olfactory sensory neurons by two anorectic peptides, insulin and leptin. J Neurophysiol 101, 2898–2906.

Simerly, R. B., Zee, M. C., Pendleton, J. W., Lubahn, D. B. and Korach, K. S. (1997). Estrogen receptor-dependent sexual differentiation of dopaminergic neurons in the preoptic region of the mouse. Proc Natl Acad Sci U S A 94, 14077–14082.

Soffker, M. and Tyler, C. R. (2012). Endocrine disrupting chemicals and sexual behaviors in fish--a critical review on effects and possible consequences. Crit Rev Toxicol 42, 653–668.

Steinfeld, R., Herb, J. T., Sprengel, R., Schaefer, A. T. and Fukunaga, I. (2015). Divergent innervation of the olfactory bulb by distinct raphe nuclei. J Comp Neurol 523, 805–813.

Stephens, S. B., Chahal, N., Munaganuru, N., Parra, R. A. and Kauffman, A. S. (2016). Estrogen Stimulation of Kiss1 Expression in the Medial Amygdala Involves Estrogen Receptor-alpha But Not Estrogen Receptor-beta. Endocrinology 157, 4021–4031.

Sundermann, E. E., Gilbert, P. E. and Murphy, C. (2008). The effect of hormone therapy on olfactory sensitivity is dependent on apolipoprotein E genotype. Horm Behav 54, 528–533.

Team, R. C. (2019). R version 3.6.1, R: A language and environment for statistical computing. R Foundation for Statistical Computing. Vienna, Austria.

Tingaud-Sequeira, A., Andre, M., Forgue, J., Barthe, C. and Babin, P. J. (2004). Expression patterns of three estrogen receptor genes during zebrafish (Danio rerio) development: evidence for high expression in neuromasts. Gene Expr Patterns 4, 561–568.

Tong, s. and Chung, B.-C. (2003). Analysis of zebrafish CYP19 promoters. The Journal of steroid biochemistry and molecular biology 86, 381–386.

Tong, S. K., Mouriec, K., Kuo, M. W., Pellegrini, E., Gueguen, M. M., Brion, F., Kah, O. and Chung, B. C. (2009). A cyp19a1b-gfp (aromatase B) transgenic zebrafish line that expresses GFP in radial glial cells. Genesis 47, 67–73.

Tyagarajan, S. K. and Fritschy, J. M. (2014). Gephyrin: a master regulator of neuronal function? Nat Rev Neurosci 15, 141–156.

Uchida, D., Yamashita, M., Kitano, T. and Iguchi, T. (2002). Oocyte apoptosis during the transition from ovary-like tissue to testes during sex differentiation of juvenile zebrafish. J Exp Biol 205, 711–718.

Valdes-Socin, H., Rubio Almanza, M., Tome Fernandez-Ladreda, M., Debray, F. G., Bours, V. and Beckers, A. (2014). Reproduction, smell, and neurodevelopmental disorders: genetic defects in different hypogonadotropic hypogonadal syndromes. Front Endocrinol (Lausanne) 5, 109.

Valverde, F., Santacana, M. and Heredia, M. (1992). Formation of an olfactory glomerulus: morphological aspects of development and organization. Neuroscience 49, 255–275.

Varshney, M. K., Inzunza, J., Lupu, D., Ganapathy, V., Antonson, P., Ruegg, J., Nalvarte, I. and Gustafsson, J. A. (2017). Role of estrogen receptor beta in neural differentiation of mouse embryonic stem cells. Proc Natl Acad Sci U S A 114, E10428–E10437.

Veyrac, A. and Bakker, J. (2011). Postnatal and adult exposure to estradiol differentially influences adult neurogenesis in the main and accessory olfactory bulb of female mice. FASEB J 25, 1048–1057.

Wang, L., Andersson, S., Warner, M. and Gustafsson, J. A. (2001). Morphological abnormalities in the brains of estrogen receptor beta knockout mice. Proc Natl Acad Sci U S A 98, 2792–2796.

Wang, L., Andersson, S., Warner, M. and Gustafsson, J. A. (2003). Estrogen receptor (ER)beta knockout mice reveal a role for ERbeta in migration of cortical neurons in the developing brain. Proc Natl Acad Sci U S A 100, 703–708.

Winter, M. J., Pinion, J., Tochwin, A., Takesono, A., Ball, J. S., Grabowski, P., Metz, J., Trznadel, M., Tse, K., Redfern, W. S., et al. Functional brain imaging in larval zebrafish for characterising the effects of seizurogenic compounds acting via a range of pharmacological mechanisms. British Journal of Pharmacology n/a.

Winter, M. J., Redfern, W. S., Hayfield, A. J., Owen, S. F., Valentin, J. P. and Hutchinson, T. H. (2008). Validation of a larval zebrafish locomotor assay for assessing the seizure liability of early-stage development drugs. J Pharmacol Toxicol Methods 57, 176–187.

Winter, M. J., Windell, D., Metz, J., Matthews, P., Pinion, J., Brown, J. T., Hetheridge, M. J., Ball, J. S., Owen, S. F., Redfern, W. S., et al. (2017). 4-dimensional functional profiling in the convulsant-treated larval zebrafish brain. Sci Rep 7, 6581.

Wullimann, M. F. (2015). Atlas of early zebrafish brain development - a tool for molecular neurogene.

Wyatt, T. D. (2003). Pheromones and animal behaviour: communication by smell and taste.

Yu, C. R., Power, J., Barnea, G., O’Donnell, S., Brown, H. E., Osborne, J., Axel, R. and Gogos, J. A. (2004). Spontaneous neural activity is required for the establishment and maintenance of the olfactory sensory map. Neuron 42, 553–566.

Zhang, L., Blackman, B. E., Schonemann, M. D., Zogovic-Kapsalis, T., Pan, X., Tagliaferri, M., Harris, H. A., Cohen, I., Pera, R. A., Mellon, S. H., et al. (2010). Estrogen receptor beta-selective agonists stimulate calcium oscillations in human and mouse embryonic stem cell-derived neurons. PLoS One 5, e11791.

